# Differential Memory Impairment Across Relational Domains in Temporal Lobe Epilepsy

**DOI:** 10.1101/2022.11.01.514752

**Authors:** Shahin Tavakol, Valeria Kebets, Jessica Royer, Qiongling Li, Hans Auer, Jordan DeKraker, Elizabeth Jefferies, Neda Bernasconi, Andrea Bernasconi, Christoph Helmstaedter, Lorenzo Caciagli, Birgit Frauscher, Jonathan Smallwood, Boris Bernhardt

## Abstract

**Background:** Temporal lobe epilepsy (TLE) is typically associated with pathology of the hippocampus, a key structure involved in relational memory processes, including episodic, semantic, and spatial memory. While it is widely accepted that TLE-associated hippocampal alterations may underlie global deficits in memory, it remains poorly understood whether TLE may present with shared or unique impairment across distinct relational memory domains.

**Methods:** We administered a recently validated behavioral paradigm to evaluate episodic, semantic, and spatial memory in 20 pharmacoresistant TLE patients and 53 age- and sex-matched healthy controls. We implemented linear mixed effects models to identify memory deficits in individuals with TLE relative to controls, and used partial least squares analysis to identify factors contributing to overall variations in relational memory performance across both cohorts.

**Results:** TLE patients showed marked impairment in episodic memory compared to controls, while spatial and semantic memory remained relatively intact. Findings were robust, with slight decreases in effect sizes after controlling for performance on executive function tests. Via partial least squares analysis, we identified group, age, and bilateral hippocampal volumes as important variables relating to relational memory impairment.

**Conclusion:** Our behavioral framework provides a granular approach for assessing relational memory deficits in people with TLE and may inform future prognostic strategies in patients with hippocampal pathology. Our work warrants further investigations into the underlying neural substrates of relational memory.

## INTRODUCTION

Temporal lobe epilepsy (TLE) is the most common pharmacoresistant epilepsy in adults, and is typically associated with pathology of the hippocampus,^1—3^ a key structure involved in declarative memory. As such, mnemonic deficits are common in TLE patients, and may sometimes impact patient quality of life more than the seizures themselves.^4,5^ A targeted investigation of behavioral phenotypes is, thus, indispensable for understanding how alterations to underlying neural substrates may affect cognition, which can inform prognostic and therapeutic approaches aimed at enhancing patient function and wellbeing.

The hippocampus supports different forms of mnemonic processes collectively termed “relational memory,” which involves the consolidation of discrete elements of subjective experience into coherent mental representations.^6—8^ Episodic, semantic, and spatial memory are specific domains of relational memory. Specifically, episodic memory integrates contiguous spatiotemporal events^9,10^ into an autobiographical abstraction known as an episode.^11,12^ In contrast, semantic memory amalgamates notions and facts into a mental hierarchy of conceptual categories.^13—15^ Finally, spatial memory maps out and binds the locations of ambient objects into a mental feature space of the physical environment, also referred to as a cognitive map.^16^ Recent studies point to a convergence of these relational domains both at the behavioral and neural level in healthy individuals.^17—25^ We have previously shown a behavioral association between semantic and spatial cognition based on performance scores obtained on different cognitive tests,^26^ which was also reflected in similar profiles of intrinsic functional connectivity between the hippocampus and neocortex.^27^ Other task-based investigations have uncovered patterns of brain activity showing six-fold symmetry, indicative of grid cell representations that capture dimensions of semantic space,^28,29^ which had only been observed in the context of physical or virtual space before.

While episodic memory impairment in TLE is well established,^1—5^ it remains unclear whether affected individuals present with difficulties in other relational memory domains. To our knowledge, integrated assessments of episodic, semantic, and spatial memory in the same participants using a standard computerized battery have not yet been conducted. Examining patients and healthy controls using a multidomain memory paradigm is, thus, an essential step in addressing the specificity of behavioral impairments across different relational dimensions resulting from TLE pathology.

In this work, we aimed to probe episodic, semantic, and spatial memory in TLE patients and healthy controls (HC) using our recently developed and open access *integrated Relational Evaluation Paradigm* (iREP, https://github.com/MICA-MNI/micaopen/task-fMRI). The iREP combines three computerized and domain-specific modules (*i*.*e*., Episodic, Semantic, and Spatial), each of which incorporates visual stimuli representing ordinary items, two levels of difficulty, and a 3-alternative forced choice design. We used linear mixed-effects models to identify behavioral associations across memory domains, levels of task difficulty, and cohorts, while controlling for underlying variations in executive function. We further implemented partial least squares analysis, a multivariate associative technique, to identify how various clinical factors contribute to shared mnemonic phenotypes across patients and controls

## MATERIALS AND METHODS

### Participants

We studied 73 adult participants, recruited between 2018 and 2022 at the Montreal Neurological Institute and Hospital, including a cohort of 20 pharmacoresistant TLE patients (9 women, mean age ± SD: 35.9 ± 11.6 years, range: 19–56, 2 ambidextrous, 12 dominant/4 non-dominant/4 unclear; see **Supplementary Material**) referred to our hospital for presurgical investigation, and 53 age- and sex-matched healthy controls (HC; 22 women, 32.1 ± 7.6 years, range: 19-57 years, 5 left-handed). Epilepsy diagnosis and seizure focus lateralization were established following a comprehensive multidisciplinary assessment based on medical history, neurological and neuropsychological evaluation, video-EEG telemetry, and MRI. Fourteen patients had a left-sided seizure focus, and 6 had a right-sided focus. Based on quantitative hippocampal MRI volumetry,^30^ 12 patients (60%) showed hippocampal atrophy ipsilateral to the focus (*i*.*e*., absolute ipsilateral-contralateral asymmetry index > 1.5 and/or ipsilateral volume z-score < -1.5). Average age at seizure onset was 22.4 ± 11.5 years (range: 2-49 years), and average duration of epilepsy was 13.5 ± 11.3 years (range: 0-38 years). All participants had normal or corrected-to-normal vision.

Our study was approved by the Research Ethics Committee of the Montreal Neurological Institute and Hospital, and all participants provided written and informed consent.

### Relational memory phenotyping

The *integrated Relational Evaluation Paradigm* (iREP) is a recently developed, open access, python-based relational memory assessment tool (https://github.com/MICA-MNI/micaopen/task-fMRI).^26^ It incorporates three complementary modules: Episodic, Semantic, and Spatial. Modules can be run either inside or outside the scanner, and are homogenized via *(i)* the use of similar visual stimuli taken from a pooled custom-made and semantically-indexed meta-library, *(ii)* the modulation of cognitive load across two conditions (*i*.*e*., Easy *vs*. Difficult) with a pseudo-randomized trial presentation order, and *(iii)* the implementation of a 3-alternative forced choice trial-by-trial paradigm. Each module contains four distinct stimulus lists (*i*.*e*., A, B, C, and D) for inter-individual counterbalancing. In the current study, all participants were tested on the iREP inside the MRI scanner, as part of a multimodal neuroimaging protocol described elsewhere.^26^ Participants used an MRI-compatible response box to provide their answers. The neural responses recorded with functional MRI are outside the scope of this study, and will be the focus of forthcoming projects.

#### (i) Episodic module

The episodic module is a symbolic version of a previously used lexicon-based episodic memory paradigm^27,31^ that involves an encoding and a retrieval run (**Fig. 1: top row**). In the encoding phase (∼6 minutes), the participant memorizes a pair of unrelated objects presented simultaneously at each trial (*i*.*e*., doorknob and ostrich). Half of the stimulus pairs is shown only once throughout the run for a total of 28 trials (*i*.*e*., Difficult condition), and the other half is displayed twice to ensure more stable encoding for a combined 56 trials (*i*.*e*., Easy condition), with a total of 84 trials for the entire task. The retrieval phase (∼4.5 minutes) is administered after a 10-min interval. During each trial, one item is displayed at the top of the monitor (*i*.*e*., doorknob) and three others, at the bottom (*i*.*e*., shark, ostrich, and ladder). From the latter three options, the participant selects the object that was paired with the top item during the encoding phase. There are 56 pseudo-randomized trials in total with equal number of trials per condition (*i*.*e*., 28 Difficult: Epi-D; 28 Easy: Epi-E).

**Figure 1.**
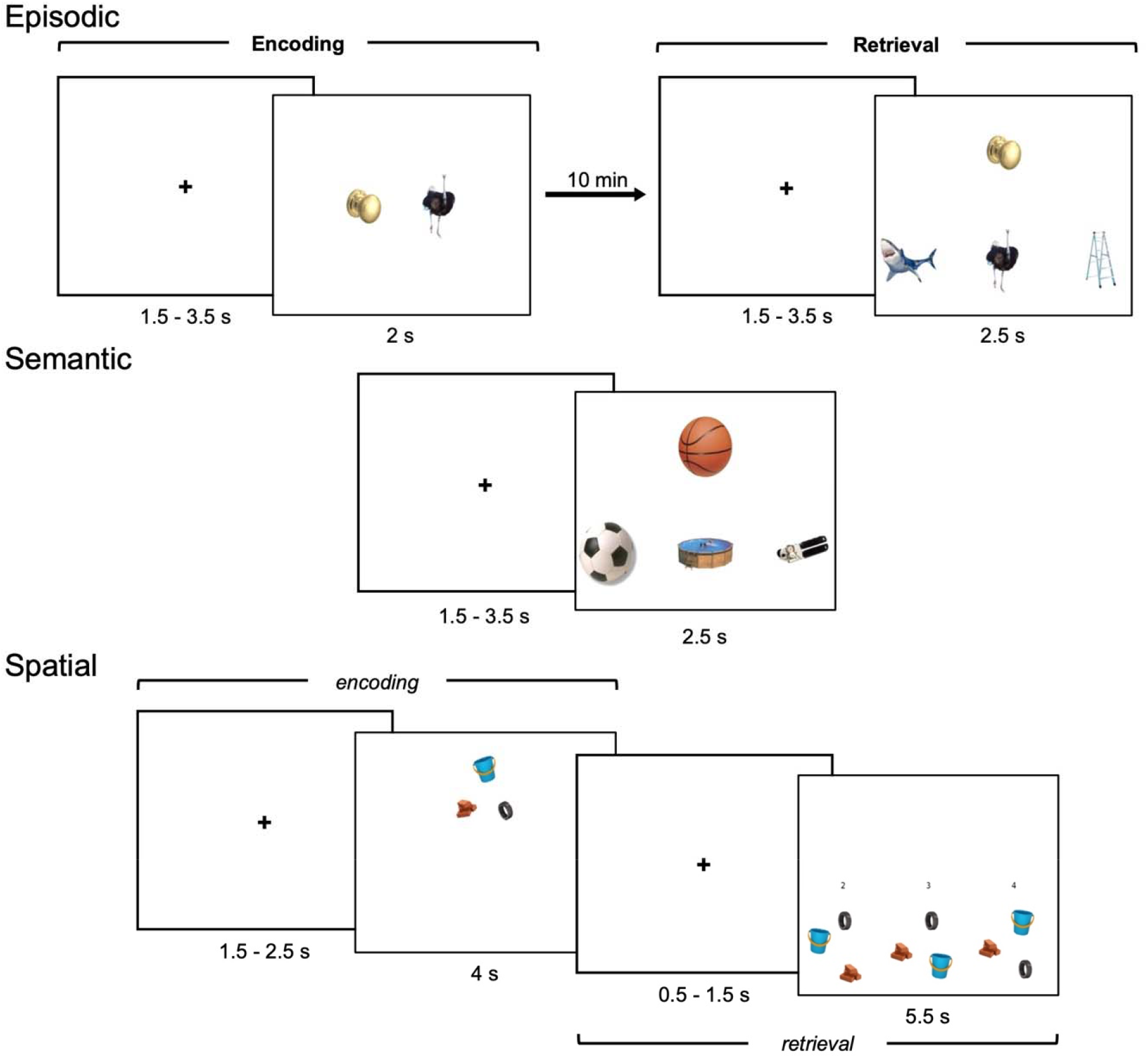
Trial design for each iREP module, *(top row)* The Episodic task consists of two separate runs. During Encoding, object pairs must be memorized. After a 10-minute break during Retrieval, the item that was originally paired with the top image must be recalled among three options, *(middle row)* In the Semantic task, the item that is the most conceptually congruent with the top object must be selected out of three choices, *(bottom row)* During the Spatial task, the configuration of three items must initially be encoded *(encoding)*. Within the same trial, the original spatial arrangement must be chosen out of three options *(retrieval)*. Numbers are there to visually aid participants on which response key to press. Overall durations for stimuli and inter-stimulus intervals are shown for each module.

#### (ii) Semantic module

The semantic module is a symbolic variant of an established lexicon-based semantic association protocol^27,32^ (**Fig. 1: middle row**). This task consists of 56 pseudo-randomized trials (∼4.5 minutes), with two conditions of equal length (*i*.*e*., 28 Difficult: Sem-D; 28 Easy: Sem-E). At each trial, a reference item appears at the top of the monitor (*i*.*e*., basketball) with three stimuli below (*i*.*e*., soccer ball, above ground pool, can opener), exactly as described in the retrieval phase of the Episodic module. The subject selects the option that is conceptually most alike to the object presented at the top. Pairwise conceptual affinity indices (*cai*) were calculated using an algorithm that leverages internet-based lexical corpora,^33^ ranging from 0 to 1. In Sem-E trials, the correct response (*i*.*e*., soccer ball) and the top image (*i*.*e*., basketball) are related by *cai > 0*.*66*; in Sem-D trials, the similarity index is given by *0*.*33 ≤ cai ≤ 0*.*66*. Regardless of condition, the conceptual relatedness of the top stimulus and the foils (*i*.*e*., above ground pool, can opener) is always *cai < 0*.*33*. Thus, the level of difficulty across conditions is a function of the semantic relationship between the top object and the correct response.

#### (iii) Spatial module

Spatial memory was assessed using a recently validated paradigm^26^ (**Fig. 1: bottom row**). In short, the Spatial module consists of 56 pseudo-randomized trials (∼12.5 minutes), with two conditions (*i*.*e*., 28 Difficult: Spa-D; 28 Easy: Spa-E). At each trial, the participant first memorizes the spatial configurations of three objects, and then selects the same arrangement among three options in a delayed-onset design. In Spa-D trials, the two distractor layouts are very similar to the target configuration as only the spacing between the objects has changed. In the Spa-E trials, in addition to the spacing, the relative position of each item within the configuration is also changed, thus making it easier to differentiate the correct arrangement from the two foils.

### Parallel assessment of executive and overall cognitive function

In addition to the iREP, we administered the EpiTrack and the Montreal Cognitive Assessment (MoCA) protocols to our participants to account for factors that could potentially confound the relationship between study cohorts and iREP outcome measures. Both tools are behavioral screening protocols for cognitive impairment. The EpiTrack is commonly used in patients with epilepsy to identify and monitor impairments in attention, processing speed, and executive function,^34,35^ while the MoCA is used to detect mild cognitive impairment and dementia.^36^

### Statistical Analysis

All data and codes used in this work are openly available at: https://github.com/MICA-MNI/micaopen/blob/master/tle_memory_manuscript_codes

#### (i) Linear mixed-effects models (LMEM)

In addition to incorporating fixed effects, LMEM also account for random effects, thus flexibly handling unequal sample sizes and multicollinearity. We implemented six different LMEM (see **Supplementary Material**) to evaluate behavioral performances as measured by percent accuracy scores, and then performed likelihood ratio tests to identify the optimal model. Specifically, this model comprised five fixed variables, including a *Group* factor with two levels (*i*.*e*., TLE, HC), a *Module* factor with three levels (*i*.*e*., Episodic, Semantic, Spatial), a *Condition* factor with two levels (*i*.*e*., Difficult, Easy), interactions *Group x Module* and *Module x Condition* terms, and a single random *Subject* term:

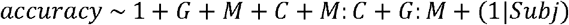

In the above formula, *accuracy* denotes percentage score. The terms *G, M*, and *C* denote group, module, and condition. The terms *M:C* and *G:M* are module-by-condition and group-by-module interactions. The term *Subj* is the random subject effect, with separate intercept to account for individual baselines. We used simple effects tests to decompose significant interactions and, where appropriate, implemented post-hoc pairwise comparisons while controlling for the false discovery rate (FDR)^37^ at α _FDR_ = 0.05.

#### (ii) Cohen’s d

To verify that significant between-group differences in relational memory performance were not driven by impairments in executive function or level of education, we computed inter-group *Cohen’s d* metrics for raw iREP accuracies and different regression models to control for: (*i*) age and sex, (*ii*) age, sex, and EpiTrack scores, (*iii*) age, sex, and MoCA scores, and *(iv)* age, sex, and education (see **Supplementary Material**). *Cohen’s d* was calculated as:

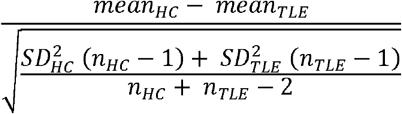

where *mean*_*HC*_, *SD*_*HC*_, and *n*_*HC*_ correspond to the average, standard deviation, and sample number for HC; *mean*_*TLE*_, *SD*_*TLE*_, and *n*_*TLE*_ are the average, standard deviation, and sample number for TLE patients.

#### (iii) Partial least squares (PLS)

PLS is a multivariate associative technique that maximizes the covariance between two datasets by decomposing their cross-correlation matrix and deriving optimal linear combinations of the original datasets known as latent components (LC).^38,39^ Unlike the factorial nature of LMEM, which seek to detect significant interactions among the various levels of predetermined variables, PLS aims to generate a lower-dimensional manifold of said factors that effectively recapitulates their raw information content. In this way, PLS offers a flexible, data-driven complementary mode of analysis. We decomposed this matrix via singular value decomposition, which resulted in a vector of left singular values (*i*.*e*., clinical saliences) characterizing a distinct clinical phenotypic pattern for each LC, a diagonal eigenvalue (*i*.*e*., singular value) matrix reflecting the covariance explained by each LC, and a vector of right singular values (*i*.*e*., iREP saliences) describing a particular iREP pattern for each LC. Subject-specific composite scores were computed by projecting their original clinical and iREP data onto their respective saliences. To test for the significance of each LC, we ran 5,000 permutation tests by resampling the iREP dataset *without* replacement while iteratively realigning permuted saliences to the original ones using Procrustes rotation to obtain a distribution of null singular values. We interpreted LCs by calculating clinical and iREP loadings, which are Pearson’s correlations between original clinical or iREP values with their corresponding composite scores (*i*.*e*., linear projections of original values onto corresponding saliences). To assess the reliability of significant LCs’ loadings, we applied a bootstrapping procedure with 5,000 iterations by resampling the iREP dataset *with* replacement and realigning bootstrapped saliences to the originals using Procrustes transform. We then estimated *z* scores for each variable loading by dividing each loading by its bootstrapped standard deviation. Finally, we converted *z* scores into FDR-adjusted *p* values (α _FDR_ = 0.05) to determine coefficient significance.

#### (iv) Exploratory analyses

We performed additional LMEM and PLS analyses in a subset of participants (LMEM: n_HC_ = 39, n_TLE_ = 16; PLS: n_HC_ = 39, n_TLE_ = 14) using weighted accuracies that incorporated reaction times: *wAcc =*(1/*RT*)\**accuracy*, where *accuracy* is the percentage score of a given module/condition, and *RT* is the average reaction time associated with that measurement.

### Hippocampal atrophy determination

We used HippUnfold^30^ to extract precise subject-specific volumes for the left and right hippocampi. HippUnfold implements a U-Net deep convolutional neural network to automate detailed hippocampal tissue segmentations. Grey matter data are then mapped onto to the resulting “unfolded” hippocampal space, with distinct subregional features. In the current work, we only examined whole hippocampal grey matter volumes, restricting analyses to MNI152-derived metrics to account for interindividual variability in intracranial volume. To compute the absolute ipsilateral-contralateral asymmetry index, we first calculated non-normalized left-right asymmetry scores for controls and patients as follows: 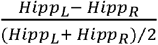, where *Hipp*_*L*_ (*Hipp*_*R*_) is the volumes of the left (right) hippocampus in MNI152 space. We normalized patient asymmetry scores with respect to those of controls, and thresholded indices at *abs(index)* > 1.5. To calculate patient ipsilateral volume z-scores, we normalized left and right volumes for patients with respect to corresponding volumes for controls, and thresholded ipsilateral values at *z*_*ipsi*_ < -1.5. Criteria for atrophy were met if either measure was satisfied (see Supplementary Material).

## RESULTS

### The structure of relational memory in HC and TLE patients: LMEM findings

Individual scores across groups and conditions are shown in **Figure 2a**. We evaluated several different LMEMs to assess iREP performance, and identified the optimal model via likelihood ratio tests (**Supplementary Table 1**), which included five fixed terms (*i*.*e*., *Group, Module, Condition, Module × Condition*, and *Group × Module*) and a random *Subject* term. Interactions were significant for *Group × Module* (F_2,348.9_ = 16.48, p < 0.001) and *Module × Condition* (F_2,345.8_ = 9.19, p < 0.001; **Supplementary Table 2**).

**Figure 2.**
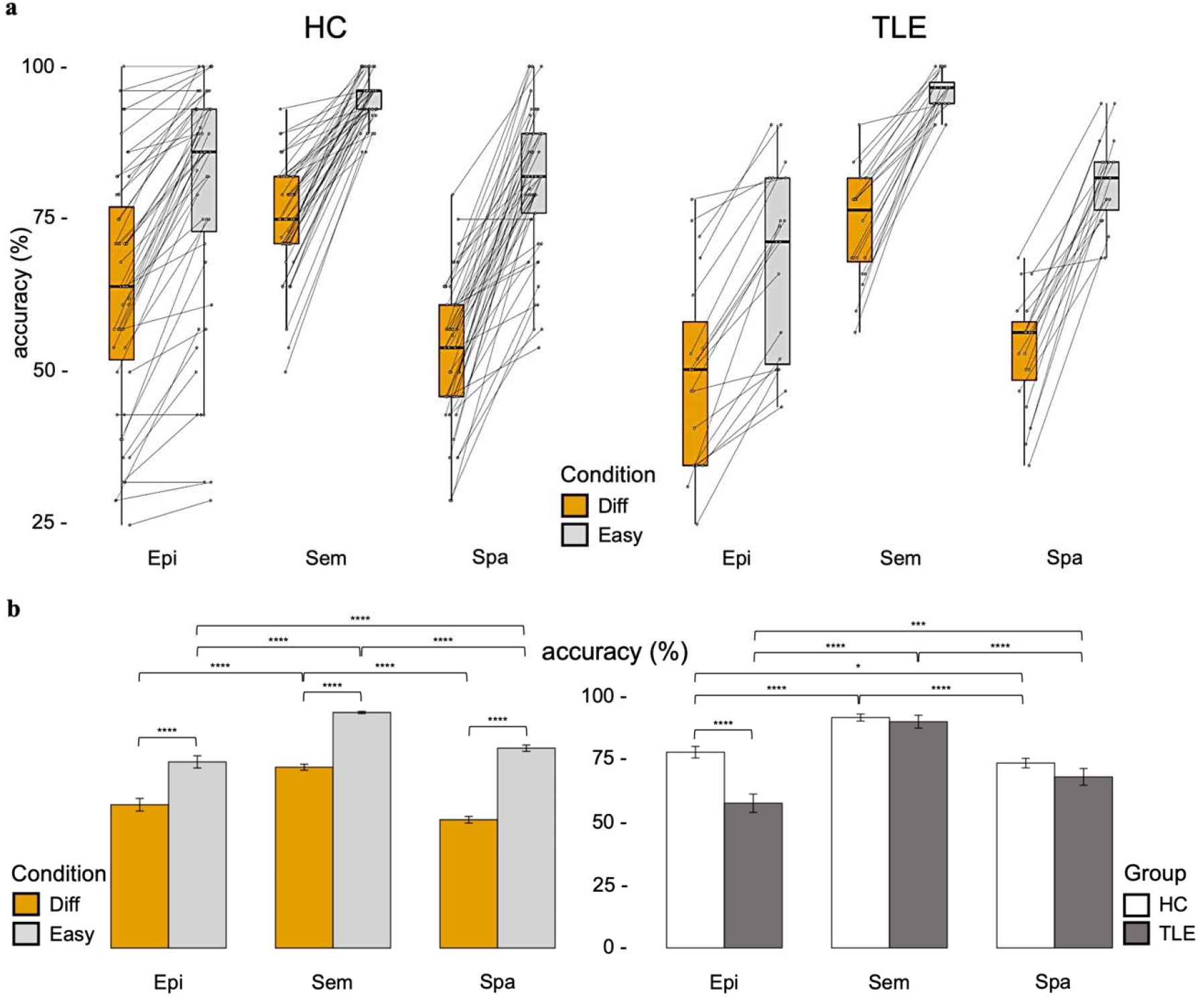
iREP performance across groups, *(a)* Performance accuracies on iREP modules show general trends across groups. Participants performed better on the easy relative to difficult condition and tended to score higher on the semantic task compared to the other two modules. Connected circles represent individual scores obtained across conditions in each task, *(b: left panel)* Module x Condition interaction. Within each module, accuracies were significantly higher on the Easy compared to Difficult condition. Semantic performance on the Difficult condition was better than either episodic or spatial. On the Easy condition, again Semantic scores were highest, followed by Spatial, and then Episodic, *(b: right panel)* Group x Module interaction. Across groups, HC outperformed TLE patients only on the Episodic task. In HC, scores were higher on the Semantic task relative to the two other modules, and Episodic accuracies were higher than in Spatial. In TLE patients, Semantic performance was also greatest, followed by Spatial, and finally Episodic (* p < 0.05, *** p < 0.001, **** p < 0.0001). Error bars represent SEM.

To decompose the *Group × Module* interaction, we conducted simple main effects tests between Groups for single iREP modules and found an effect of *Group* on Episodic performance only (Episodic: F_1,68_ = 14.00, p < 0.001; Semantic: F_1,141_ = 0.36, p = 0.550; Spatial: F_1,142_ = 2.16, p = 0.144). FDR-adjusted pairwise comparisons confirmed lower performance on this module in TLE compared to controls (t_174_ = 6.40, p < 0.0001, **Figure 2b: right panel**). Additional main effects tests between modules for each Group revealed a strong *Module* effect on accuracies, irrespective of cohort (controls: F_2,256.19_ = 27.42, p < 0.0001; TLE: F_2,95.95_ = 27.76, p < 0.0001). Here, controls scored highest on the Semantic module, outperforming Episodic accuracies (t_349_ = 8.09, p < 0.0001), and scored higher on the Episodic relative to the Spatial module (t_350_ = 2.38, p < 0.05). Similarly, TLE patients scored higher on the Semantic relative to the Spatial module (t_350_ = 7.92, p < 0.0001). Unlike controls, however, TLE patients obtained higher scores on the Spatial relative to the Episodic module (t_353_ = 3.47, p < 0.001, **Figure 2b: right panel**).

We also decomposed the *Module × Condition* interaction to illustrate how performance on the modules differed irrespective of group (Episodic: F_1,69.00_ = 169.91, p < 0.0001; Semantic: F_1,60.28_ = 336.88, p < 0.0001; Spatial: F_1,71.00_ = 401.62, p < 0.0001). As expected, performances were consistently higher on the Easy relative to Difficult condition (ts ≥ 8.92, ps < 0.0001, **Figure 2b: left panel**). We observed additional main effects between modules for each condition (Difficult: F_2,140.21_ = 51.62, p < 0.0001; Easy: F_2,138.25_ = 46.39, p < 0.0001). Pairwise tests indicated that in the Difficult condition, Semantic scores were higher than either Episodic or Spatial module scores (t scores _≥_ 9.37, ps < 0.0001). Semantic accuracies in the Easy condition were, similarly, greater than either Episodic or Spatial (t scores _≥_ 7.42, ps < 0.0001), and scores were higher on the Spatial compared to the Episodic module (t_350_ = 4.12, p < 0.0001, **Figure 2b: left panel**).

The *Group x Condition* interaction was not captured by this model, but did show a trend towards significance in other sub-optimal models (M3: F_1,348.3_ = 2.75, p = 0.098; M5: F_1,342.5_ = 3.31, p = 0.070, see **Supplementary Material**). These findings were supported by between-group *Cohen’s d* statistics calculated for raw iREP scores (Epi-E: 0.73, Epi-D: 1.00, Sem-E: -0.22, Sem-D: 0.37, Spa-E: 0.43, Spa-D: 0.67), suggesting differences between the two cohorts that were more marked on the Difficult relative to Easy condition across modules (see **Supplementary Table 4**).

When accounting for reaction times, exploratory LMEM analyses with weighted accuracies further extended these results. Participants scored highest on the Easy relative to Difficult condition across tasks, and Semantic performance was superior to both Episodic and Spatial. Interestingly, both groups scored lowest on the Spatial task (**Supplementary Figure 1, left panel & Supplementary Tables 3.1, 3.2, 3.4**), and while controls outperformed TLE patients on the Episodic module once again, they outscored them on Spatial as well (**Supplementary Figure 1, right panel & Supplementary Tables 3.1, 3.3, 3.5**), suggesting that these weighted accuracies may have been more sensitive to spatial memory deficits in TLE.

### The structure of relational memory in individuals with TLE and controls: PLS findings

To complement LMEM findings, we implemented PLS to ascertain the presence of a clinical profile associated with iREP scores, and found that age, group, and hippocampal volume contributed to relational memory performance. The first latent component (LC1) obtained via the decomposition of the cross-correlation matrix between clinical phenotypes and iREP accuracies accounted for more than 81% of the total covariance (**Figure 3a, left**). The correlation between corresponding clinical and behavioral composite scores along LC1 was also highly significant, as attested by permutation tests of its singular value (r = 0.46, p_perm_ < 0.001, **Figure 3a, right**). We also ran an additional bootstrapping scheme to evaluate the robustness of LC1 loadings (age: - 0.64, sex: -0.17, group: 0.83, left/right hippocampal volumes: 0.54/0.63, Epi-E: 0.82, Epi-D: 0.85, Sem-E: 0.25, Sem-D: 0.49, Spa-E: 0.55, Spa-D: 0.65, **Figure 3b, left**). Not including sex (*z* = -1.42, p = 0.193), all clinical and iREP variables presented with significantly reliable loadings on LC1 (age: *z* = -7.31, group: *z* = 23.29, left/right hippocampal volumes: *z =* 4.88/5.80, Epi-E: 25.56, Epi-D: 22.05, Sem-E: 2.37, Sem-D: 4.73, Spa-E: 5.69, Spa-D: 8.61, all p_FDR_ < 0.05, **Figure 3b, right)**.

**Figure 3.**
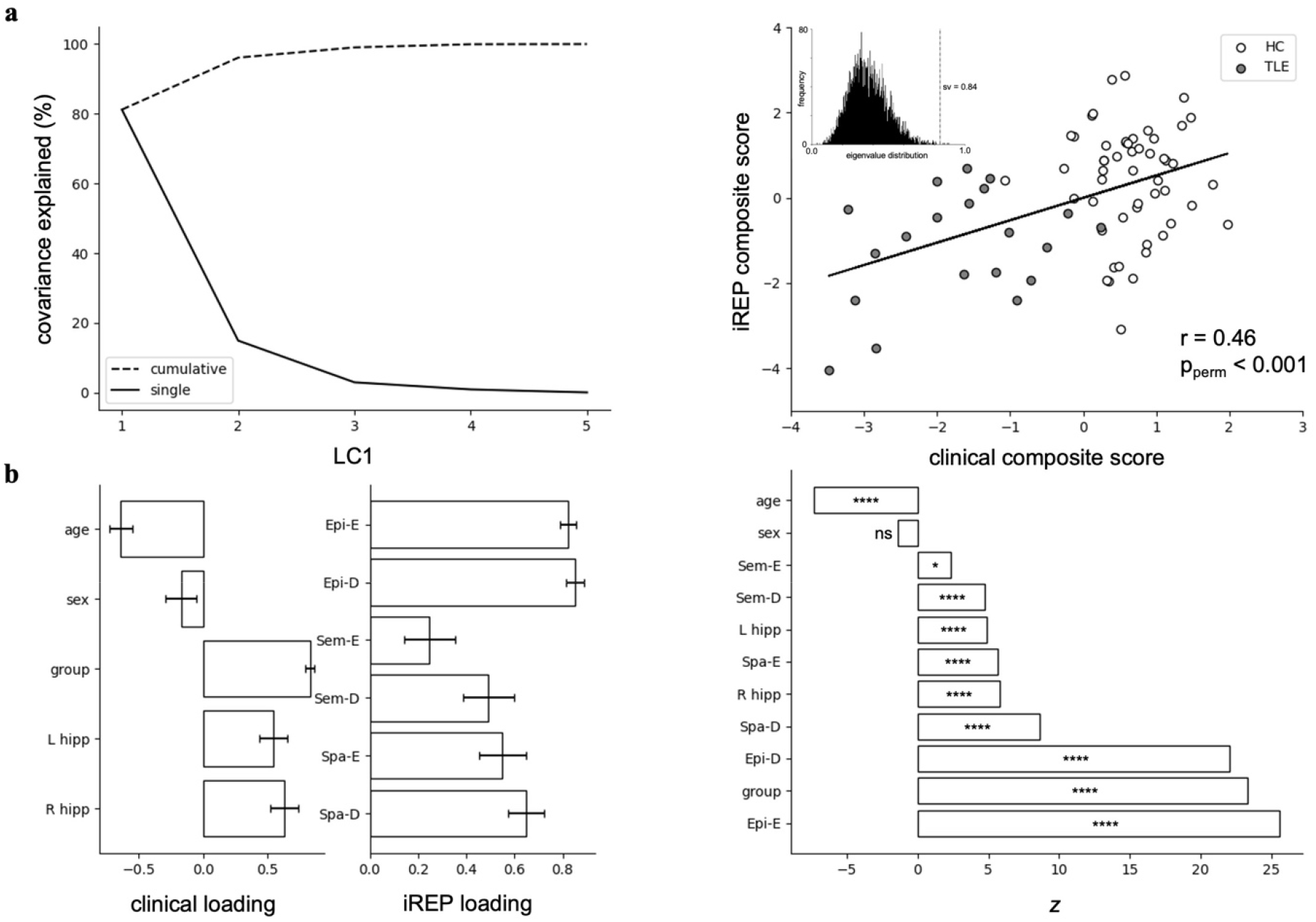
PLS. *(a) left:* the first latent component (LC1) accounted for over 81% of the covariance between five clinical features *(i*.*e*., age, sex, group, and left/right hippocampal volumes) and six iREP measurements *(i*.*e*., Epi-E, Epi-D, Sem-E, Sem-D, Spa-E, and Spa-D). *right:* the association between clinical and iREP composite scores along LC1 was significant (r = 0.46, p_penn_ < 0.001) as attested by 5,000 permutations (inset: dashed line “sv” represents the actual singular value), *(b) left:* clinical and iREP loadings with standard deviations (error bars), *right:* loading reliabilities were determined by estimating z scores for each variable by dividing loading coefficients by standard deviations derived from 5,000 bootstraps. *Z* scores were adjusted for FDR (* p_FDR_ < 0.05, **** p_FDR_ < 0.0001).

Thus, younger age, allocation to the HC cohort, and larger bilateral hippocampal volumes were associated with better performance across all task measurements, and while the iREP pattern was shared across modules, episodic accuracies showed highest contributions, followed by spatial, and finally semantic, validating LMEM findings. Overall, diagnostic group and episodic scores were the most important features of LC1. These findings were further validated in exploratory PLS analyses with weighted accuracies that incorporated reaction times (**Supplementary Figure 2**).

### Assessment of executive function and general cognitive impairment

We probed associations between the above LMEM and PLS results and more general differences in cognitive and executive function. As our LMEM findings showed a significant difference in Episodic scores between TLE patients and controls, we repeated the between-group comparisons after regressing out EpiTrack and MoCA scores separately, and computed *Cohen’s d* values for each regression model. We found that the effects of the group differences in Episodic outcomes across conditions were overall reduced by as much as 38.7% in Epi-E and 29.1% in Epi-D, but nonetheless remained moderate-to-high (Epi-E: *Cohen’s d* > 0.45; Epi-D: *Cohen’s d* > 0.71, **Supplementary Table 4**). We also correlated iREP composite scores for LC1 with measurements obtained on the EpiTrack and MoCA. In both cases we observed moderate correlations (EpiTrack: r = 0.36, p < 0.01; MoCA: r = 0.40, p < 0.001), suggesting that these variables have a shared, yet not identical effect.

## DISCUSSION

Our objective was to analyze the differential behavioral impairments across separate relational memory domains in patients with temporal epilepsy, the most common pharmacoresistant epilepsy in adults and a human disease model of memory dysfunction. We compared the performances of TLE patients to those of age- and sex-matched healthy controls on the different modules of the *integrated Relational Memory Paradigm* (iREP), our recently developed cognitive assessment tool. The iREP is a comprehensive mnemonic protocol that includes three complementary and homogenous tasks that collectively tap into the episodic, semantic, and spatial processing systems of the brain. Modules are further stratified into two conditions that correspond to levels of difficulty, thus offering two degrees of probing resolution into each cognitive domain. We applied linear mixed effects models (LMEM) in conjunction with partial least squares (PLS) analysis to identify general associations in behavioral scores across groups, modules, and conditions, and to discern latent associative patterns between clinical features and performance scores.

Our LMEM results confirmed that TLE patients were considerably impaired on the episodic module, a finding that expands on an already well-established scientific corpus.^1—5^ Also, PLS analysis revealed that group allocation and performance scores on both conditions of the episodic task were the strongest contributors to the first PLS latent component, further validating the notion of episodic deficits in TLE. Moreover, we deciphered additional contributions from left and right hippocampus volumes, supporting a potential link between the integrity of the hippocampi and relational cognition in general, and episodic memory specifically. Hippocampal contributions to relational memory performance are well established, and its decline in TLE is related to many factors, including subregional structural pathology,^40^ disruptions in connectivity patterns,^41^ and functional reorganization.^42^ Overall, our PLS findings confirmed associations between clinical presentation and general mnemonic ability, pointing specifically to episodic impairments in TLE patients that exacerbate as a function of increasing age and decreasing hippocampal volumes. We were also interested in whether more general impairment in cognitive and executive function, attention, and processing speed might have contributed to the observed between-group differences in episodic memory.^43,44^ Thus, we administered supplemental behavioral screening tools to ensure that group disparities were not driven solely by neurobehavioral differences in other domains. Specifically, we used the EpiTrack and MoCA,^34— 36^ which are designed to track deficits in executive function and attention as well as mild cognitive impairment and dementia, respectively. While performances on the EpiTrack and MoCA correlated with PLS-derived iREP composite scores, group differences in episodic memory persisted even after controlling for these screening tests, suggesting that these differences were not uniquely mediated by non-relational cognitive domains.

Interestingly, peak scores in both cohorts were achieved on the Semantic module, where TLE patients performed on par with controls. While the Episodic and Spatial tasks encompass built-in phases for stimulus encoding and retrieval, the Semantic consists of retrieval only. Presumably, the underlying conceptual associations between objects required to complete this module successfully were incidentally and repeatedly encoded throughout the participant’s lifetime, implicating long-term memory consolidation, which benefits not only from hippocampal but also non-hippocampal neocortical contributions.^45^ Indeed, insofar as TLE patients present with semantic deficits, faulty encoding of novel conceptual relations has been suggested as a potential cause.^46^ This consideration is in line with the complementary learning systems framework, which posits a division of labour underlying memory and learning, whereby the hippocampus rapidly encodes non-overlapping episodic representations that are gradually consolidated into a latent semantic structure across the neocortex through interleaved reinstatement of episodic engrams.^47,48^ Likewise, the multiple trace theory stipulates a resilience of the semantic memory system to lesions of the hippocampus, a structure, which, in contrast to its recurrent involvement in binding disparate neocortical patterns that code for either episodic or spatial information, is surmised to be only transiently active in the context of semantic cognition.^49^ In addition, we also note that semantic impairments in people with TLE are typically measured using visual confrontation naming tasks like the Boston Naming Test, which, while suitable for identifying dysnomia, do not necessarily tap into semantic association processes per se.^50,51^ In fact, TLE patients seem to be relatively intact on semantic assessment protocols similar to our own where conceptual judgment is required,^51,52^ such as the Intelligenz-Struktur-Test, where an outlier must be selected out of five lexical alternatives (*i*.*e*., sitting, lying, *going*, kneeling, standing).^53^ While research is ongoing to elucidate the network dynamics involved in verbal deficiencies associated with TLE,^54^ behavioral divergence across verbal and non-verbal domains may offer an avenue for mapping out phenotypic differences between TLE and other similar neurological conditions, such as semantic dementia, in which patients appear to be impaired on both domains.^55^ Even though the Semantic module of the iREP is a valid test of general conceptual knowledge,^26,27,32^ the absence of group differences in the current work does not necessarily entail that TLE patients are unaffected on more sensitive measures of semantic cognition, as it has been shown that impairments may emerge if tasks are sufficiently difficult.^56^ Notwithstanding more fine-grained forms of conceptual processing, we can conclude that memory of general associations between everyday items is relatively well retained in TLE.

Patients scored marginally lower than controls on the spatial task, but results were not statistically significant. It is known that the hippocampus supports allocentric spatial memory, a mode of spatial processing that involves the three-dimensional relations between objects in an environment independent of the subjective viewpoint,^57—60^ with the volume of the hippocampus further associated with proficiency in this allocentric domain.^61—63^ Therefore, we were expecting to see clear indications of spatial deficits in the TLE cohort given that performance on the spatial module was previously shown to be correlated with the Four Mountains Task,^26^ an established protocol for examining allocentric spatial memory in clinical populations that present with hippocampal pathology.^57,64^ However, it has been reported that allocentric spatial cognition might be generally well preserved in patients who present with mild hippocampal sclerosis, short disorder duration, and low seizure frequency, even in right-sided lesions typically associated with cognitive impairment in this domain.^65^ Moreover, deficits in individuals with medial temporal lobe lesions scale with the magnitude of the probing delay between stimulus encoding and retrieval, with relatively intact short-term memory of spatial information for short delays.^43,66^ As alluded to, visuo-spatial and figural memory impairments are predominantly observed in patients who present with a right-sided seizure focus,^67—72^ which only accounted for 30% of our TLE cohort. Consequently, the combined effect of relatively short inter-trial probing delays (0.5s - 1.5s) with a comparatively small sub-sample of right-sided TLE patients may have contributed to downplaying the impact of individual spatial deficiencies at the group level. In forthcoming studies, we will be increase sample sizes to test for latent impairments in allocentric spatial cognition. We should also note that when incorporating reaction times in our analyses, sensitive increased to show additional between-group differences on the Spatial task, which was further captured in the latent pattern of association between clinical features and iREP scores with greater contributions from Spatial measurements. From a design perspective, while iREP modules are all predicated on a 3-alternative forced choice design, the Spatial task stands out from the rest given the complexity of its stimuli. Whereas in the other two modules, the three choices during retrieval are each composed of a single object (*i*.*e*., one object is one choice), in the spatial task, each choice is in fact three separate items that combine to make a triangular configuration of objects. We posit that this added layer of stimulus complexity may have translated into relatively longer reaction times, which impacted Spatial weighted accuracies. While weighted measurements that incorporate both percentage scores and reaction times offer a relatively comprehensive summary of behavioral outcomes, they may overinflate the variance in the data and skew statistical inferences as a result,^73^ which is why we implemented them for purely exploratory purposes.

Collectively, our findings demonstrate atypical behavioural patterns of relational memory in TLE patients. In particular, they underscore marked impairments in episodic memory (for item pairs with no semantic cues) associated with age and hippocampal volume. On the other hand, memory for conceptual associations appeared preserved, and there were signs of subtle alterations in the efficiency of spatial memory. Given stringent diagnostic criteria for inclusion in our TLE cohort, which resulted in a relatively small sample of 20 pharmacoresistant patients, we had to make some unavoidable compromises. While we acknowledge that seizure onset, seizure laterality, and hemispheric dominance are important factors that can affect behavioral outcomes, we omitted these variables from our study because of sample size constraints. We continue to expand our patient cohort and hope to account for these factors in future work. Even so, our initial observations already provide detailed insights into the differential impairment across relational memory domains accompanying hippocampal damage in TLE, warranting complementary investigations into underlying neural substrates. As in previous efforts,^26^ we will characterize the morphological and functional and connectome level correlates of relational memory processes as indexed by the iREP in future work. We are hopeful that this novel mnemonic paradigm will be positively received by the neuroscientific community and actively implemented to address neurobehavioral variations in memory function in health and disease.

## SUPPLEMENTARY MATERIAL

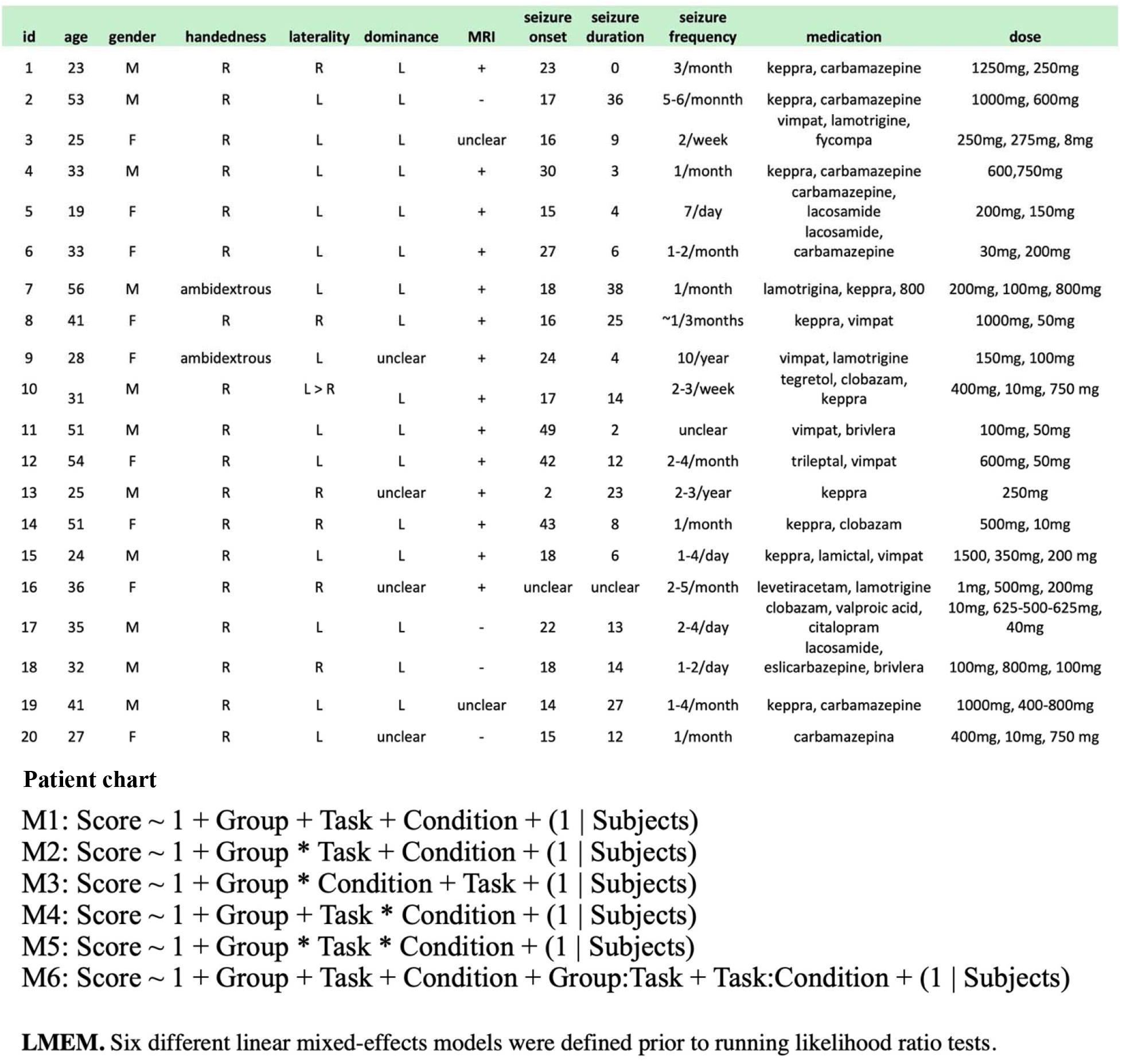

**Table 1.** Likelihood ratio tests. M6 was the optimal model for fitting the behavioral scores with linear mixed-effects. (n.par: number of parameters; A1C: Akaike’s Information Criteria; BIC: Bayesian Information Criteria; logLik: log-likelihood; **** p < 0.0001)

| Model | n.par | AIC | BIC | logLik | deviance | $\chi^2$ | df | p |
| --- | --- | --- | --- | --- | --- | --- | --- | --- |
| M1 | 7 | 3435.1 | 3463.5 | -1710.6 | 3421.1 | -- | -- | -- |
| M3 | 8 | 3434.4 | 3466.8 | -1709.2 | 3418.4 | 2.7725 | 1 | 0.0959 |
| M2 | 9 | 3408.5 | 3445.0 | -1695.3 | 3390.5 | 27.8206 | 1 | 1.331e-07 **** |
| M4 | 9 | 3422.4 | 3459.0 | -1702.2 | 3404.4 | 0.0000 | 0 | -- |
| M6 | 11 | 3394.3 | 3438.9 | -1686.1 | 3372.3 | 32.1589 | 2 | 1.039e-07 **** |
| M5 | 14 | 3396.9 | 3453.7 | -1684.4 | 3368.9 | 3.4164 | 3 | 0.3318 |

**Table 2.** Linear mixed-effects model. Significant interaction effects were observed for Module x Condition and Group x Module. (SS: sum of squares; MS: mean squares; Num/Den df: numerator/denominator df)

| Variable | SS | MS | Num df | Den df | F | p |
| --- | --- | --- | --- | --- | --- | --- |
| Group | 1880 | 1880 | 1 | 69.44 | 14.3158 | 0.0003244 *** |
| Module | 30803 | 15401 | 2 | 348.92 | 117.2694 | < 2.2e-16 **** |
| Condition | 54663 | 54663 | 1 | 345.77 | 416.2169 | < 2.2e-16 **** |
| Module:Condition | 2413 | 1206 | 2 | 345.77 | 9.1855 | 0.0001298 *** |
| Group:Module | 4328 | 2164 | 2 | 348.92 | 16.4764 | 1.453e-07 **** |

**Table 3.1.** Exploratory LMEM with weighted accuracies. Significant interaction effects were observed for Module x Condition and Group x Module. (SS: sum of squares; MS: mean squares; Num/Den df: numerator/denominator df)

| Variable | SS | MS | Num df | Den df | F | p |
| --- | --- | --- | --- | --- | --- | --- |
| Group | 2471 | 2471 | 1 | 54.54 | 23.4927 | 0.00001081 **** |
| Module | 64076 | 32038 | 2 | 265.43 | 304.6341 | < 2.2e-16 **** |
| Condition | 30072 | 30072 | 1 | 261.16 | 285.9380 | < 2.2e-16 **** |
| Module:Condition | 4717 | 2358 | 2 | 261.15 | 22.4243 | 1.029e-09 **** |
| Group:Module | 1355 | 677 | 2 | 265.43 | 6.4418 | 0.001854 ** |

**Table 3.2.** Simple main effects for Module x Condition. Analysis of variance conducted with Satterthwaite’s method.

| Interaction | SS | MS | Num df | Den df | F | p |
| --- | --- | --- | --- | --- | --- | --- |
| Condition @ Episodic | 6904.4 | 6904.4 | 1 | 53 | 104.32 | 3.998e-14 **** |
| Condition @ Semantic | 22667 | 22667 | 1 | 49.644 | 694.7 | < 2.2e-16 **** |
| Condition @ Spatial | 3981.5 | 3981.5 | 1 | 54 | 185.43 | < 2.2e-16 **** |
| Module @ Difficult | 25970 | 12985 | 2 | 105.5 | 127.68 | < 2.2e-16 **** |
| Module @ Easy | 61458 | 30729 | 2 | 104.95 | 205.57 | < 2.2e-16 **** |

**Table 3.3.** Simple main effects for Group x Module. Analysis of variance conducted with Satterthwaite’s method.

| Interaction | SS | MS | Num df | Den df | F | p |
| --- | --- | --- | --- | --- | --- | --- |
| Group @ Episodic | 2160.3 | 2160.3 | 1 | 52 | 11.204 | 0.001523 ** |
| Group @ Semantic | 1120 | 1120 | 1 | 102 | 3.5835 | 0.06119 |
| Group @ Spatial | 869.41 | 869.41 | 1 | 108 | 12.996 | 0.000474 *** |
| Module @ HC | 61403 | 30702 | 2 | 191.54 | 116.1 | < 2.2e-16 **** |
| Module @ TLE | 21970 | 10985 | 2 | 87 | 75.015 | < 2.2e-16 **** |

**Table 3.4.** Post-hoc pairwise comparisons for Module x Condition. P-values adjusted for FDR.

| Module | contrast | estimate | SE | df | t | p |
| --- | --- | --- | --- | --- | --- | --- |
| Episodic | Diff - Easy | -16 | 1.97 | 260 | -8.103 | < 0.0001 **** |
| Semantic | Diff - Easy | -30 | 2.01 | 261 | -14.896 | < 0.0001 **** |
| Spatial | Diff - Easy | -12 | 1.96 | 260 | -6.153 | < 0.0001 **** |
| Condition | contrast | estimate | SE | df | t | p |
| Difficult | Episodic - Semantic | -11.8 | 2.12 | 262 | -5.559 | < 0.0001 **** |
| Difficult | Episodic - Spatial | 18.1 | 2.08 | 263 | 8.705 | < 0.0001 **** |
| Difficult | Semantic - Spatial | 29.9 | 2.10 | 264 | 14.195 | < 0.0001 **** |
| Easy | Episodic - Semantic | -25.8 | 2.13 | 262 | -12.101 | < 0.0001 **** |
| Easy | Episodic - Spatial | 22.1 | 2.08 | 263 | 10.609 | < 0.0001 **** |
| Easy | Semantic - Spatial | 47.8 | 2.12 | 264 | 22.586 | < 0.0001 **** |

**Table 3.5.**
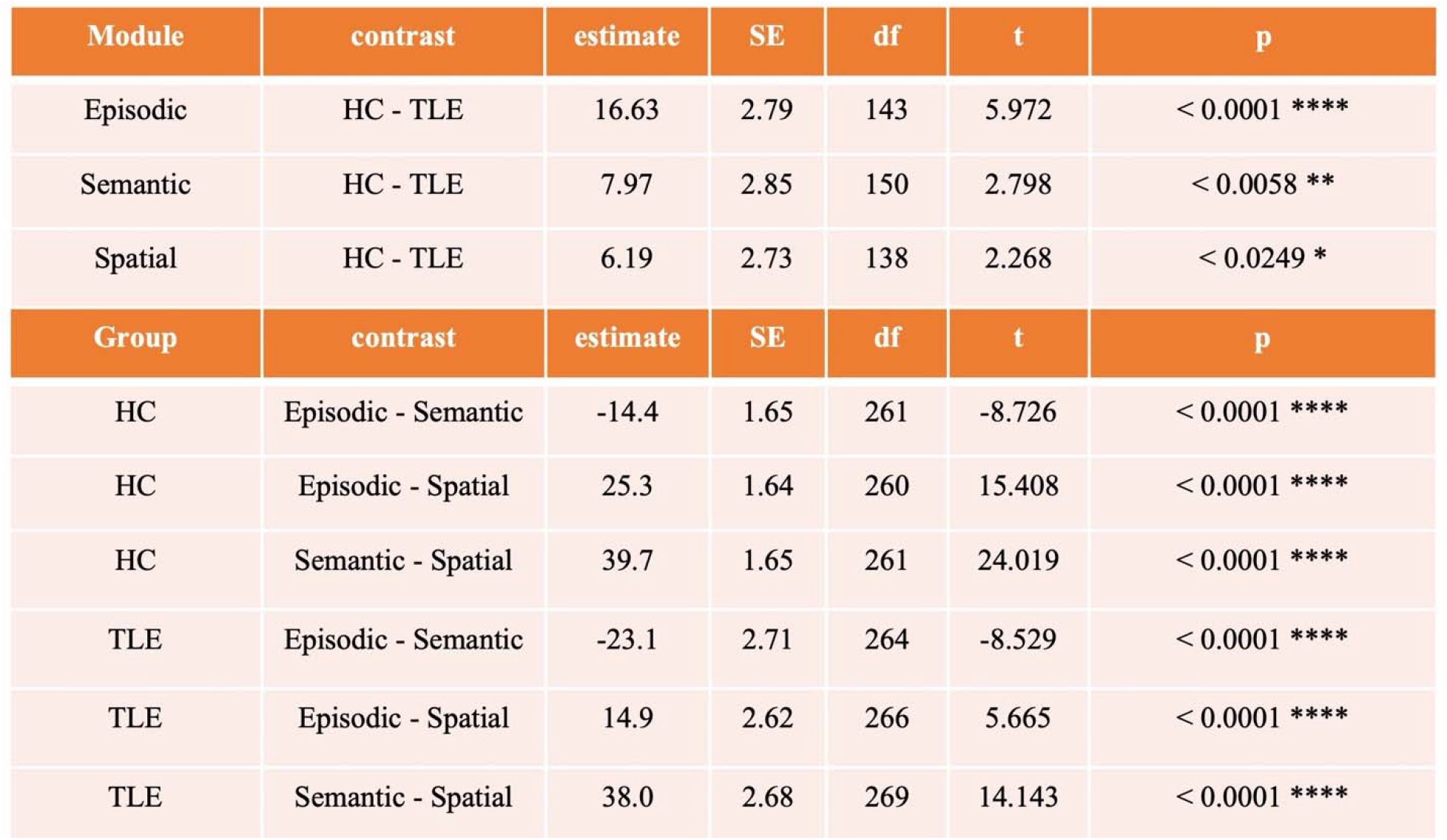
Post-hoc pairwise comparisons for Group x Module. P-values adjusted for FDR.

**Table 4.** Cohen’s d. Values depict raw, age/sex (AS) controlled, age/sex/EpiTrack (ASE) controlled, age/sex/MoCA (ASM) controlled, and aged/sex/education (ASEdu) controlled inter-group effect sizes. Level of education was quantified as: 1 for high school, 2 for undergraduate, and 3 for graduate.

| measure | Cohen's d | nHC | nTLE |
| --- | --- | --- | --- |
| raw Epi-E | 0.7270 | 53 | 20 |
| raw Epi-D | 1.0010 | 53 | 20 |
| raw Sem-E | -0.2185 | 53 | 20 |
| raw Sem-D | 0.3727 | 53 | 20 |
| raw Spa-E | 0.4302 | 53 | 20 |
| raw Spa-D | 0.6744 | 53 | 20 |
| AS Epi-E | 0.6523 | 53 | 20 |
| AS Epi-D | 0.9099 | 53 | 20 |
| AS Sem-E | -0.2321 | 53 | 20 |
| AS Sem-D | 0.3493 | 53 | 20 |
| AS Spa-E | 0.3009 | 53 | 20 |
| AS Spa-D | 0.5812 | 53 | 20 |
| ASE Epi-E | 0.4455 | 51 | 15 |
| ASE Epi-D | 0.7097 | 51 | 15 |
| ASE Sem-E | -0.2814 | 51 | 15 |
| ASE Sem-D | 0.3784 | 51 | 15 |
| ASE Spa-E | 0.2126 | 51 | 15 |
| ASE Spa-D | 0.6324 | 51 | 15 |
| ASM Epi-E | 0.6328 | 52 | 20 |
| ASM Epi-D | 0.8912 | 52 | 20 |
| ASM Sem-E | -0.2470 | 52 | 20 |
| ASM Sem-D | 0.3283 | 52 | 20 |
| ASM Spa-E | 0.2748 | 52 | 20 |
| ASM Spa-D | 0.5601 | 52 | 20 |
| ASEdu Epi-E | 0.6770 | 49 | 20 |
| ASEdu Epi-D | 1.0020 | 49 | 20 |
| ASEdu Sem-E | -0.1845 | 49 | 20 |
| ASEdu Sem-D | 0.3893 | 49 | 20 |
| ASEdu Spa-E | 0.3513 | 49 | 20 |
| ASEdu Spa-D | 0.5837 | 49 | 20 |

**Figure 1.**
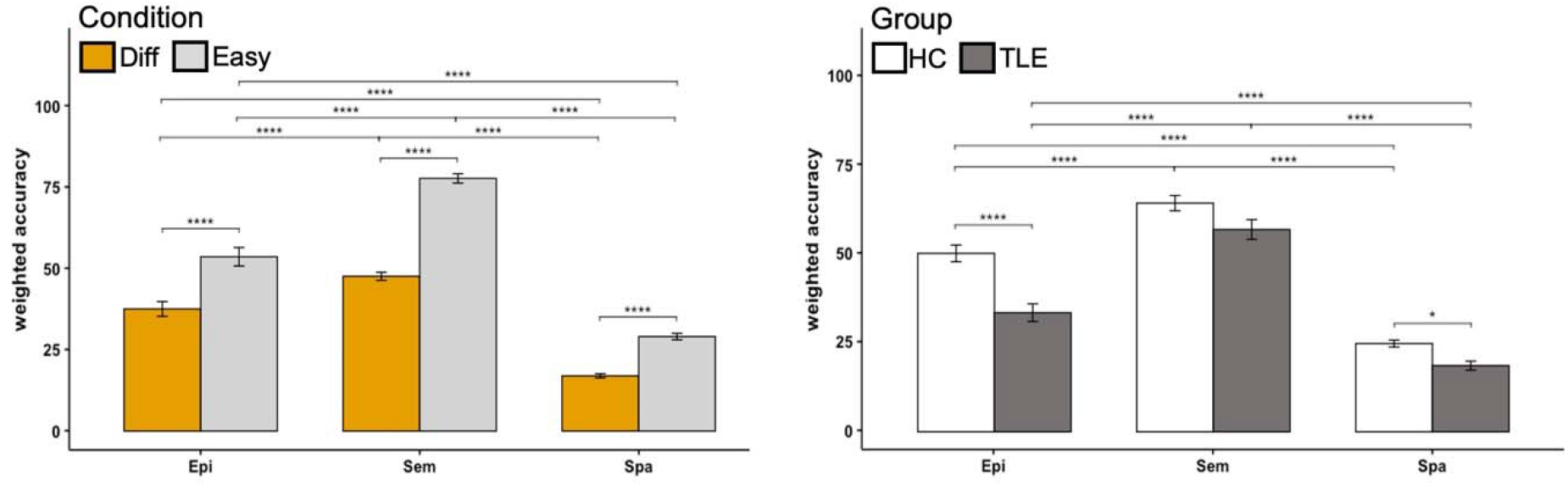
LMEM with weighted accuracies. Average reaction times were extracted for 55 participants (nHC = 39, nTLE = 16) across module/condition to generate weighted accuracies *(i*.*e*., 1/RT * accuracy), which were then incorporated into exploratory LMEM analyses. Findings recapitulated prior observations, but also indicated that participants were overall more affected on the spatial module relative to episodic or semantic (left panel), with additional group-wise differences on the spatial module (right panel, * p < 0.05, **** p < 0.0001).

**Figure 2.**
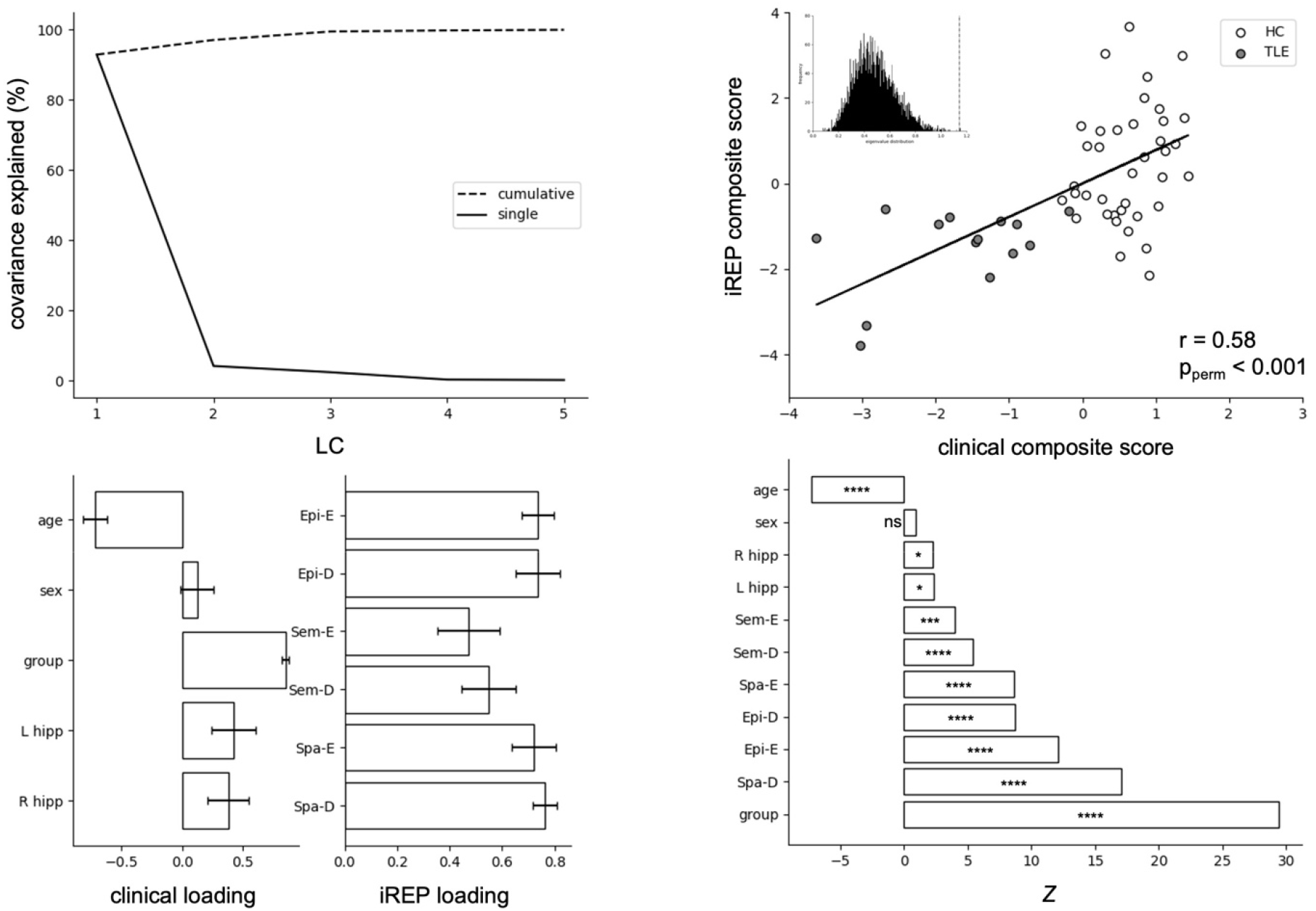
PLS with weighted accuracies. PLS findings for 53 participants (nHC = 39, nTLE = 14) corroborate previous results, indicating that LC1 accounts for over 92% of the covariance (top left), with a high degree of correspondence between clinical and iREP composite scores (r = 0.58, p_perm_ < 0.001, top right) as attested by 5,000 permutations (inset), and loading patterns (bottom left) that validate associations between diagnostic group, age, hippocampal volume and iREP performance, with greater contribution from Spa-D performance (bottom right). Estimated *z* scores were adjusted for FDR (* prDR < 0.05, *** p_FDR_ < 0.001, **** p_FDR_ < 0.0001).

